# Unleashing natural competence in *Lactococcus lactis* by induction of the competence regulator ComX

**DOI:** 10.1101/147132

**Authors:** Joyce Mulder, Michiel Wels, Oscar P. Kuipers, Michiel Kleerebezem, Peter A. Bron

## Abstract

In biotechnological work horses like *Streptococcus thermophilus* and *Bacillus subtilis* natural competence can be induced, which facilitates genetic manipulation of these microbes. However, in strains of the important dairy starter *Lactococcus lactis* natural competence has not been established to date. However, *in silico* analysis of complete genome sequences of 43 *L. lactis* strains revealed complete late-competence gene-sets in 2 *L. lactis* subspecies *cremoris* strains (KW2 and KW10) and 8 *L. lactis* subspecies *lactis* strains, including the model strain IL1403 and the plant-derived strain KF147. The remainder of the strains, including all dairy isolates, displayed genomic decay in one or more of the late competence genes. Nisin-controlled expression of the competence regulator *comX* in *L. lactis* subsp. *lactis* KF147 resulted in the induction of expression of the canonical competence regulon, and elicited a state of natural competence in this strain. By contrast, *comX* expression in *L. lactis* NZ9000, predicted to encode an incomplete competence gene-set, failed to induce natural competence. Moreover, mutagenesis of the *comEA-EC* operon in strain KF147, abolished the *comX* driven natural competence, underpinning the involvement of the competence machinery. Finally, introduction of nisin-inducible *comX* expression into *nisRK*-harboring derivatives of strains IL1403 and KW2 allowed the induction of natural competence also in these strains, expanding this phenotype to other *L. lactis* strains of both subspecies.

**Significance statement:** Specific bacterial species are able to enter a state of natural competence in which DNA is taken up from the environment, allowing the introduction of novel traits. Strains of the species *Lactococcus lactis* are very important starter cultures for the fermentation of milk in the cheese production process, where these bacteria contribute to the flavor and texture of the end-product. The activation of natural competence in this industrially relevant organism can accelerate research aiming to understand industrially relevant traits of these bacteria, and can facilitate engineering strategies to harness the natural biodiversity of the species in optimized starter strains.

## Introduction

Horizontal gene transfer (HGT) fulfills an important role in the evolution of bacteria (1–4). In several species, an important mechanism for HGT is natural competence. This phenomenon is defined as a cellular state that enables internalization of exogenous DNA, followed by autonomous replication as a plasmid or incorporation into the chromosome via homologous recombination. Among Gram-positive bacteria, natural competence was first described in *Streptococcus pneumoniae* (5, 6). More recently, it was found that among lactic acid bacteria (LAB), the important yoghurt bacterium *Streptococcus thermophilus* can enter a state of natural competence upon culturing in chemically defined medium (7). When Gram-positive bacteria enter a state of natural competence, exogenous DNA translocates through the DNA-uptake machinery, a multiprotein complex comprising the proteins ComEA, ComEC, ComFA, ComFC and a nuclease (EndA in *S. pneumoniae*) encoded by the late competence (*com*) genes (8, 9). Other late competence genes encode for proteins that compose pili-like structures (ComGA-GG) or protect internalized DNA against degradation (SsbA, SsbB, DprA and RecA) (8, 9). Expression of these genes is positively regulated by the competence master-regulator ComX, that acts as an alternative sigma factor (10–12). In *S. thermophilus*, expression of *comX* is initiated upon formation of the quorum sensing ComRS complex comprising the pheromone-like peptide ComS and transcriptional regulator ComR, encoded by the *comRS* operon (13, 14). Addition of a synthetic peptide that resembles the active competence pheromone has proven a successful strategy to induce natural competence in several bacterial species. For example, addition of a synthetic ComS peptide to *S. thermophilus* cultures in the early logarithmic phase of growth enabled the activation of natural competence and highly efficient DNA transformation (13, 15). Analogously, other streptococci including *S. pneumoniae* utilize the *comCDE* regulatory module to control natural competence, involving the competence-stimulating peptide (CSP, encoded by *comC*) and a two-component system (encoded by *comD* and *comE*, (16, 17)), and the addition of synthetic CSP leads to development of natural competence in this species.

Strains of *Lactococcus lactis* are of great importance in the dairy industry, primarily in the production of cheese and butter(milk) (18). So far, a *comRS*- or *comCDE*-like system has not been identified in *L. lactis*. Nevertheless, complete sets of late competence genes appear to be present in several *L. lactis* genomes (19, 20, 21 and this study). In addition, increased expression of competence genes has been observed in *L. lactis* subsp. *lactis* IL1403 and KF147 under specific conditions that included carbon starvation (22, 23). Unfortunately, in neither of these strains, or any other *L. lactis* strain, natural competence development could be experimentally established (19, 23). As an alternative route to establish natural competence, overexpression of *comX* has been employed, aiming to enhance expression of the complete late competence regulon. Such an approach has been successful in *S. thermophilus* (24), but failed in *L. lactis* IL1403 (19). Nevertheless, the observation that complete sets of late competence genes are apparently present in some of the *L. lactis* genomes (25, 26) and that their expression can be induced under specific conditions (22, 23), deserves a more dedicated bioinformatic and experimental effort.

Here, we present a comparative genomics analysis of 43 *L. lactis* genomes to assess their potential to enter a state of natural competence. Moreover, by employment of controlled expression of ComX we demonstrate enhanced expression of the late competence regulon and concomitant induction of natural competence, which was only successful in strains predicted to encode a complete late competence machinery. The discovery of natural competence in *L. lactis* will enable transfer of genetic information in a non-GMO manner, resulting in the improvement of the industrial performance of strains of this species and the enhancement of fermented product quality.

## Results

### Genomic analyses show complete sets of competence genes in several *L. lactis* strains

To evaluate whether *L. lactis* strains possess the genetic capacity to enter a state of natural competence, late-competence associated genes were initially identified in the *L. lactis* KF147 genome by using the known competence genes of the naturally competent *Streptococcus thermophilus* LMD-9 (7, 13). This strain was selected for this primary analysis based on previous work that reported that many of the late competence genes were induced in this strain under starvation, non-growing conditions (23). Similar proteins (both in length and sequence) were identified to be encoded within the KF147 genome for all selected late competence genes of *S. thermophilus* LMD-9, corroborating that a complete competence gene-set is present in this *L. lactis* strain (Table S2).

Subsequently, the identified KF147 competence protein sequences were used as a reference set for identification and comparison to the orthologous groups (OGs) of genes encoded by 42 other *L. lactis* strains (20, 21, 25–34). Full length protein sequence identity relative to the KF147 OGs was calculated for all 42 *L. lactis* strains (Fig. 1). This analysis revealed considerable genomic decay in several of the strains of both the *lactis* and *cremoris* subspecies. Moreover, for the OGs that were intact, there was a clear distinction between the levels of identity observed for strains belonging to the subspecies *lactis* (that includes strain KF147) and *cremoris* (Fig. 1), exemplifying the genetic distinction between these two subspecies (35). Among the strains belonging to the *cremoris* subspecies, only strains KW2 and KW10 appeared to encode full length homologues of all the late competence proteins selected in KF147, albeit with identity scores ranging from 56 to 99% (Fig. 1). Notably, when the late competence gene-set of the KW2 strain was used to determine full-length protein sequence identity levels, instead of those of strain KF147, it was apparent that late competence proteins displayed a high degree of conservation within the *cremoris* subspecies, but were distinct from their orthologues in the *lactis* subspecies (Fig. S1). Among the 28 strains belonging to the subspecies *lactis*, 9 appeared to encode a full set of late competence proteins.

**Fig. 1:**
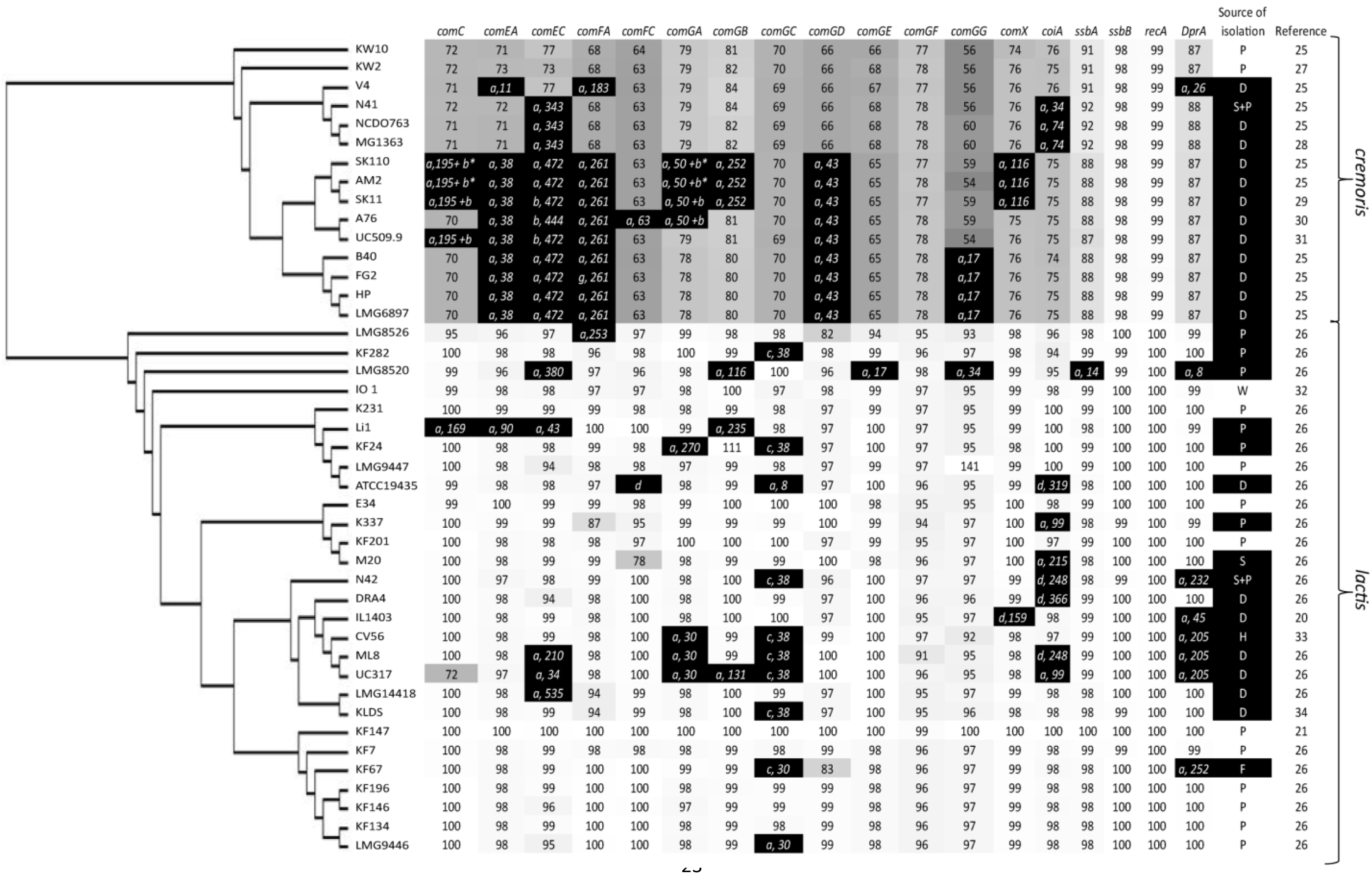
Genomic analysis of 43 *L. lactis* strains to assess genetic capacity to develop natural competence. A concatenated core genome SNP tree of 43 *L. lactis* strain was combined with full-length protein identity-scores (%) for the selected subset of late competence associated proteins in comparison to their homologues in strain *L. lactis* KF147 that was used as a reference. Protein identity scores are depicted within each cell and reflected by gray scales based on the *L. lactis* KF147 query protein sequences in which 90%< full length alignments are considered as presence of the full length of the competence gene. Genetic events leading to competence gene decay (black cells in the figure) are specified as premature stopcodon within the first 90% of the gene (a), transposon insertion (b), prophage insertion (c) absence of gene, mutated/alternative start or lengthened/fused protein; at least more than 25% of its total length (d), followed by the position within the protein sequence where the event is detected relative to its N-terminus. The source of isolation column letters represent: P= plant, D= dairy, S= soil, W= water, H= human body and F= fruit (69) and references for the genome sequences when available.

The genomic decay within these late competence genes in the subspecies *cremoris* strains displayed several conserved disruptive mutations in specific genes, including IS*982* insertions in *comEC* (strains SK11, A76 and UC509.9) and *comGA* (strains SK11, and A76), although with some variation with respect to the precise position of insertion (Fig. 1 and Fig. S2). Various strains of the *cremoris* subspecies contained conserved premature stop codons within one or more of their late competence genes, suggesting that these strains derive from a common ancestor, in which conserved and strain specific mutations have shaped the decay pattern of the late competence genes. For example, strains SK110, AM2, SK11, A76, UC5099, B40, FG2, HP and LMG6897 share similar mutational events in *comEA, comEC, comFA* and *comGD*, whereas strains N41, NCDO763 and MG1363 harbor common mutations in *comEC* (Fig. S3). In contrast, the disruptive mutations observed in the late competence genes of strains of the subspecies *lactis* appeared more scattered (Fig. 1), suggesting that degenerative mutations accumulated more recently in this subspecies. Nevertheless, several strains (KF282, KF24, N42, CV56, ML8, KLDS, UC317, KF67) contain a (remnant of a) prophage insertion within the *comGC* gene (Fig. S4). Remarkably, these phage sequences are always inserted at the same position within the *comGC* sequence, suggesting site-specific integration at a conserved sequence element within the *comGC* gene.

In summary, these findings indicate that in the majority of *L. lactis* strains one or more late competence functions are compromised, suggesting that these strains are not able to develop a state of natural competence. The analysis also implies that in some strains, including *L. lactis* KF147, the genetic capacity to enter a state of naturally competence appears to be intact. Finally, it is noteworthy that within the present panel of strains, there are no dairy isolates that appear to encode a complete set of intact late competence proteins, which may reflect the high-level of genome decay that has been reported for strains in the milk environment before (36–38).

### Moderate overexpression of the late competence regulon regulator ComX results in a state of natural competence in *L. lactis* KF147

In order to test whether the identified competence machinery can be activated and is functional, we set out to overexpress the predicted competence regulator ComX. From the subset of strains predicted to harbor a complete set of competence genes, *L. lactis* KF147 harbors a chromosomal copy of *nisRK* but does not produce nisin (39), allowing nisin-inducible *comX* expression by cloning of this gene under control of P_*nisA*_ in pNZ8150 ((40). This *comX* expression strategy led to a dose-dependent inhibition of growth (Fig. 2A), which was not observed in the control strains harboring pNZ8150 (Fig. S5; (40)), or pNZ8040, a vector enabling nisin-inducible expression of *pepN* (Fig. S5; (41)). Hence, the observed growth retardation is not caused by the addition of nisin or the overexpression of proteins as such but is specifically caused by the presence of ComX.

**Fig. 2:**
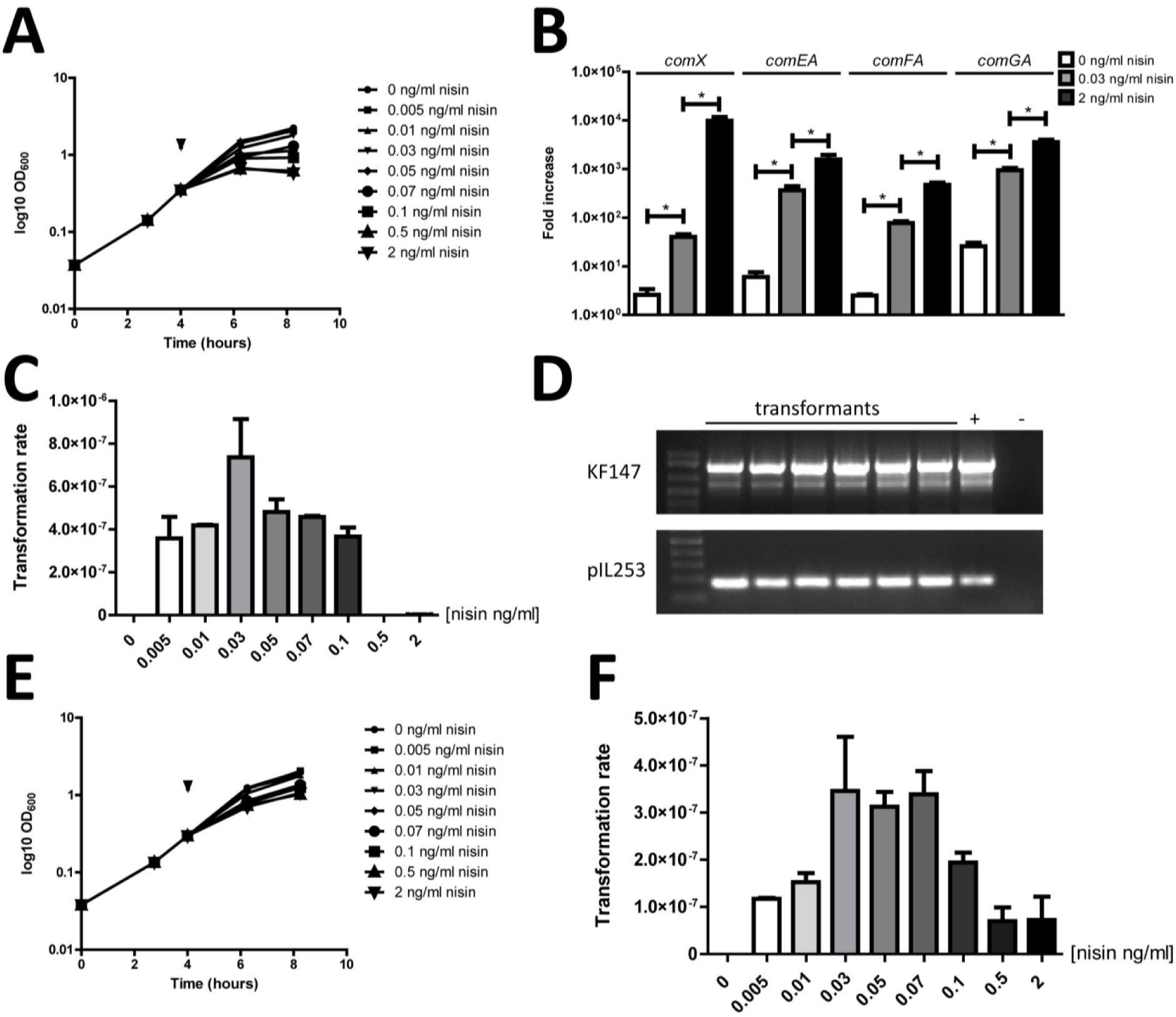
Impact of ComX expression on growth, late competence gene expression and natural competence phenotype in *L. lactis* KF147. Dose dependent growth inhibition upon nisin induction of *L. lactis* KF147 harboring pNZ6200 (panel A). The arrow indicated the time point of nisin induction. Panel B represents *comX, comEA, comFA* and *comGA* expression levels after nisin induction, whereas an asterix indicates significant differences (P<0.05). Number of colonies obtained (panel C) and confirmation of their genetic identity are presented in panels C and D, respectively. Panel E and F display the same analysis in strain *L. lactis* KF147 harboring pNZ6201.

To investigate the impact of elevated ComX levels on the expression level of the late competence genes, their transcript levels were determined by RT-qPCR on RNA derived from *L. lactis* KF147 harboring pNZ8150 or pNZ6200, either uninduced, or moderately or fully induced with nisin. In uninduced conditions, *comX* expression levels were 2.5- to 6-fold increased in *L. lactis* KF147 harboring pNZ6200 as compared to the pNZ8150 harboring cells, which is likely reflecting low-level ‘promoter leakage’ due to the presence of P_*nisA*_ on a high-copy plasmid (Fig. 2B). Induction of *comX* expression in *L. lactis* KF147 harboring pNZ6200 with either 0.03 or 2 ng/ml nisin for 2 hours led to 15-20 and 1500-4000-fold induction of *comX* expression relative to the uninduced control of the same strain, respectively (Fig. 2B). Similarly, expression of the late competence genes *comEA, comFA*, and *comGA* was induced, illustrating the strongly enhanced expression of the late competence regulon as a consequence of the elevated levels of its regulator ComX (Fig. 2B). These induction conditions for the activation of late competence genes were employed to test whether the corresponding phenotype could also be observed, by adding pIL253 to the culture medium at the same timepoint that *comX* induction was initiated using a range of nisin concentrations. As expected, no pIL253 transformants were obtained for *L. lactis* KF147 harboring pNZ8150 under any of the conditions tested (data not shown). By contrast, pIL253 transformants were obtained for *L. lactis* KF147 harboring pNZ6200 following induction with nisin concentrations ranging from 0.005 to 0.1 ng/ml nisin, with an approximate transformation rate of 10^−7^−10^−6^ (transformants / total cell number μg plasmid DNA). The highest transformation rates were obtained after 0.03 ng/ml nisin induction (Fig. 2C). Both strain identity and pIL253 presence was confirmed by PCR in all transformants tested (Fig. 2D). Notably, full nisin-induction (2.0 ng/ml nisin) of *comX* expression in pNZ6200 harboring *L.lactis* KF147 did not result in any transformants. To check whether *comX* of the *cremoris* strain MG1363 is still functional, transformation of *L. lactis* KF147 harboring pNZ6201 upon nisin induction was also tested. Similar results were obtained when *comX* of *L. lactis* subsp. *cremoris* MG1363 was expressed in *L. lactis* subsp. *lactis* KF147 (Fig. 2E, F), indicating that *comX* derived from a *cremoris* strain is also fully functional. Taken together, these results demonstrate that activation of moderate expression, but not high-level expression, of endogenous *comX* in *L. lactis* KF147 elicits the natural competence phenotype in this strain. The observation that this does not occur at a high level of *comX* expression may be a consequence of the observed growth defect under these conditions, which may interfere with completion of the competence machinery assembly and/or recovery of potential transformants after plating. Such notion, is supported by the observation that expression of a heterologous copy of *comX* (derived from *L. lactis* subspecies *cremoris*) induced less severe growth defects upon maximal nisin induction, and still led to detectable natural competence development, albeit with reduced efficiency as compared to moderate nisin induction levels.

### ComX-induced transformation in *L. lactis* depends on the late competence operon *comEA-EC*

The experiments above do not provide direct proof for a functional dependency of the observed transformation phenotype on the expression of the late competence genes, although this is likely, considering the fact that these genes encode the DNA-uptake machinery. Therefore, we constructed a *comEA-EC* negative derivative of *L. lactis* KF147 through the integration of a linear fragment harboring a tetracyclin-resistance encoding *tetR* flanked by regions homologous to the 5’- and 3’- regions surrounding the *comEA-EC* operon. The procedure for moderate *comX* induction was applied to transform this linear mutagenesis fragment to strain KF147 harboring pNZ6200. Integrants with the anticipated genotype (Δ*comEA-EC::tetR;* Fig. 3A) were obtained with a similar efficiency as was observed for pIL253 transformation. Subsequent *comX* expression induction experiments in the Δ*comEA-EC::tetR* derivative of strain KF147 (NZ6200) harboring pNZ6200. Full induction of *comX* expression led to a growth rate reduction in this strain (Fig. 3B) similar to what was observed for the parental KF147 strain. Importantly, nisin concentration-dependent *comX* overexpression and corresponding upregulation of expression of *comFA* and *comGA*, but not *comEA*, genes was similar to what was established in *L. lactis* KF147 harboring pNZ6200 (Fig. S6). However, in contrast to the parental strain KF147, transformation of NZ6200 harboring pNZ6200 with pIL253 did not yield any transformants (Fig. 3C), establishing the involvement of the *comEA-EC* operon in *comX*-induced competence in *L. lactis* KF147.

**Fig. 3:**
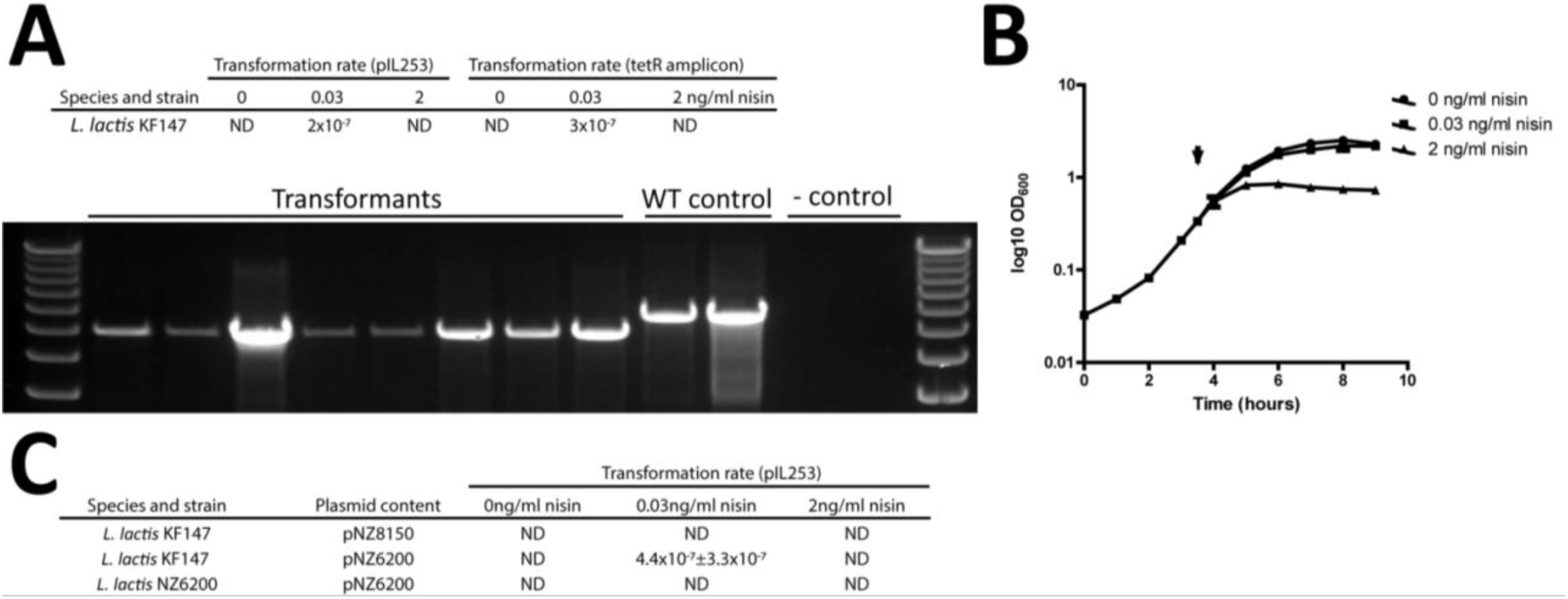
Natural competence of *L. lactis* KF147 depends on the late competence operon *comEA-EC.* By employment of a linear mutagenesis fragment, a *comEA-EC* negative derivative of KF147 harboring pNZ6200 (NZ6200) could be obtained with high efficiency (panel A). NZ6200 displayed a similar growth defect upon full nisin induction (panelB) but transformation with pIL253 is not feasible (panel C).

### Expansion of the natural competence phenotype to a broader set of *L. lactis* strains

In order to employ the comparative genomics analysis performed in this study as a predictor for competence potential, *L. lactis* subspecies *cremoris* strains KW2 (27), and NZ9000 (40, 42), and *L. lactis* subspecies *lactis* IL9000 (43) were tested for transformability upon *comX* induction. Strains NZ9000 and IL9000 are derivatives of MG1363 and IL1403, respectively, which contain the *nisRK* genes integrated in their chromosome, thereby allowing the use of the nisin controlled expression system. Strain KW2 does not harbor *nisRK* in its genome and to facilitate the use of the nisin induction system in this strain, pNZ9531 was first introduced into this strain, thereby expressing *nisRK* in this background from a low-copy plasmid vector that is compatible with the pNZ8150 backbone of the pNZ6200 and pNZ6201 vectors used for *comX* expression (44). The plasmids pNZ8150, pNZ6200 or pNZ6201 were transformed to electrocompetent cells of IL9000, NZ9000, and pNZ9531-harboring *L. lactis* KW2. These transformants were induced with 0, 0.03 or 2 ng/ml nisin, and the induction of the competence phenotype was evaluated in these strains by transformation with pIL253 (for NZ9000 and IL9000) or pNZ6202 (a tetracycline-selectable pIL253 derivative). As anticipated, none of the conditions employed allowed the activation of natural competence in strain NZ9000 (Table 1), which is in good agreement with the *comEC* and *coiA* mutations observed in its parental strain MG1363 (Fig. 1). In contrast, transformants were obtained for pNZ6200 and pNZ6201 harboring derivatives of IL9000, and when these plasmids were transformed to KW2 harboring pNZ9531 (Table 1). Notably, although the efficiency of transformation appeared to be the highest for the cells in which moderate *comX* expression was induced (i.e., 0.03 ng/ml), also under uninduced and upon high level induction of *comX* expression (i.e., 2 ng/ml) transformants were obtained. The observed transformation under uninduced conditions might be caused by the previously reported higher levels of ‘leakage’ of the *nisA* promoter activity in *L. lactis* strains harboring the *nisRK* expression vector pNZ9531 (44), whereas differential expression of *nisRK* and/or the *dprA* mutation in the IL9000 parental strain (IL1403; Fig. 1) may require lower levels of *comX* expression to activate competence, as the DprA function has been associated with competence shut-off (45, 46). The observation that high-level *comX* expression still allowed competence development in KW2 and IL9000 despite the growth-inhibitory consequences of this level of induction, which is in contrast with the results obtained for strain KF147, remains to be determined. Finally, in pNZ9531-harboring KW2, induced expression of the *comX* derived from *L. lactis* subspecies *cremoris* (i.e., as expressed from pNZ6201) allowed competence development, confirming the bidirectional functional exchangeability of the *comX* genes from these two *L. lactis* subspecies. These results confirm the predictions made by comparative genomics (Fig. 1) with respect to the capacity to develop a natural competence phenotype in *L. lactis* strains, and establish that strains of both subspecies have the capacity of natural competence which can be induced by controlled expression of the *comX* encoded regulator from either of the subspecies.

**Table 1:** Assessing transformation with plasmid pIL253/pNZ6202 by controlled expression of *comX* in *L. lactis* IL9000, NZ9000, and pNZ9531 harboring KW2. Transformation rate is calculated as number of pIL253 or pN6202 transformants/total number of cells/μg DNA, ND= not detected.

| Species and strain | Plasmid content | Transformation rate (pIL253/pNZ6202) |  |  |
| --- | --- | --- | --- | --- |
|  |  | 0ng/ml nisin | 0.03ng/ml nisin | 2ng/ml nisin |
| *L. lactis* IL9000 | pNZ6200 | $1.5 \times 10^{-7} \pm 8.2 \times 10^{-8}$ | $1.7 \times 10^{-7} \pm 1.5 \times 10^{-7}$ | $3.4 \times 10^{-7} \pm 1.7 \times 10^{-7}$ |
| *L. lactis* IL9000 | pNZ8150 | ND | ND | ND |
| *L. lactis* KW2 | pNZ9531+ pNZ6200 | $1.5 \times 10^{-6} \pm 1.1 \times 10^{-6}$ | $1.0 \times 10^{-5} \pm 9.2 \times 10^{-7}$ | $5.2 \times 10^{-6} \pm 3.1 \times 10^{-6}$ |
| *L. lactis* KW2 | pNZ9531+ pNZ6201 | $1.1 \times 10^{-6} \pm 1.1 \times 10^{-6}$ | $6.2 \times 10^{-6} \pm 5.7 \times 10^{-6}$ | $3.1 \times 10^{-7} \pm 2.2 \times 10^{-7}$ |
| *L. lactis* KW2 | pNZ9531+ pNZ8150 | ND | ND | ND |
| *L. lactis* NZ9000 | pNZ6200 | ND | ND | ND |
| *L. lactis* NZ9000 | pNZ8150 | ND | ND | ND |

## Discussion

This study demonstrates that the *L. lactis* strains KF147, KW2 and IL1403 possess a functional DNA-uptake machinery, which can be activated by the ComX regulator. This implies that identification of a complete set of late competence genes through comparative genomics represents an appropriate approach to predict the capacity of a strain to enter a state of natural competence, and it seems likely that most, if not all, of the other strains identified here to encode complete gene sets can be made naturally competent via the same strategy of *comX* overexpression. It should be noted that the expression of a much larger set of over 100 genes is regulated upon addition of the competence pheromone in streptococci (11, 47, 48). For instance, development of natural competence usually occurs in concert with increased expression of proteins involved in DNA recombination, thereby facilitating integration of acquired DNA (49), a feature that has been observed in this study for *L. lactis* KF147 as well suggesting expression of such proteins in *L. lactis* KF147 upon competence development. Nevertheless, we show that the dedicated assessment of only the canonical late competence genes is a valid predictor for competence potential in *L. lactis* strains.

It is commonly assumed that the *L. lactis* ancestor strain prior to subspeciation into the *lactis* and *cremoris* subspecies originates from a plant-associated niche and that strains adapted to increase their fitness in the nutritionally rich dairy environment (39, 50, 51). Remarkably, none of the dairy isolates of *L. lactis* that were analyzed here appears to encode a complete set of late-competence genes, suggesting that during the adaptation to the dairy niche there is no significant environmental fitness benefit associated with the capacity to become naturally competent. This may relate to a real lack of fitness benefit of this phenotype within the dairy environment or may be due to highly consistent suppression of the phenotype during growth in milk, thereby preventing the possible fitness benefit to become apparent, which may allow the decay of encoding genes without apparent fitness cost for the bacteria. The latter scenario appears to be in agreement with the observed activation of the expression of late competence genes in *L. lactis* upon carbon starvation conditions (22, 23), that are not likely to occur within the dairy niche, as it is very rich in lactose. The genomic decay events associated with dairy-derived *L. lactis* strains include prophage disruption of the *comGC* locus in strains of the subspecies *lactis* (27, 52), and insertion of IS*982* into several *com* genes in strains of the *cremoris* subspecies (39, 53–55). Notably, the phylogenetic relatedness of *L. lactis* subspecies *cremoris* strains predicted on basis of competence gene-decay events, displayed a topology that was remarkably similar to that observed for the core-genome relatedness of these strains (Fig. S3). Importantly, typical dairy environment associated lactic acid bacteria quite commonly display genomic decay as a consequence of the adaptation to this nutritionally rich environment (36–38). For example, loss of function events have been observed in *S. thermophilus, Lactobacillus helveticus* and *Lactobacillus bulgaricus* upon prolonged culturing in milk, with mutations accumulating in genes encoding transport, energy metabolism and virulence associated functions, implying that these functions do not contribute to fitness in the dairy niche (36, 38, 56, 57). Analogously, experimental evolution of *L. lactis* KF147 to enhanced fitness and growth in milk was shown to be associated with suppression of gene repertoires associated with the import and utilization of a variety of typically plant-environment associated carbon sources, as well as mutations leading to functional reconstitution and elevated transcription of the peptide import system (*opp*) of this strain [52]. Paradoxically, dairy strains of *S. thermophilus* still possess the genetic and phenotypic capacity to develop natural competence (13, 24, 38, 58), suggesting that competence development in this species contributes to fitness in this habitat. Contrary to *L. lactis*, where carbon starvation has been associated with induction of late competence expression (22, 23), similar conditions have not been implied in competence regulation in *S. thermophilus*. This may suggest that *S. thermophilus* actively expresses the competence phenotype in the dairy environment, which may contribute to this species’ fitness in the milk environment.

In nature, natural competence in bacteria is commonly a transient phenotypic state with a small window of opportunity to take up DNA (59), of which activation and shut-down are subject to subtle regulation (45, 46) to prevent futile activation of the costly process and to sustain genomic stability. Analogously, optimal induction in *L. lactis* was achieved with a moderate level of ComX induction, whereas high-level induction of this regulator failed to lead to competence development (strain KF147), or led to significantly reduced levels of transformation (strains KW2, and IL9000). Moreover, high level *comX* expression was consistently associated with reduced growth efficiency of the strains used in this study, which is illustrative of the tight connection between competence and growth (60). Previous *comX* expression studies using *L. lactis* IL1403, the parental strain of IL9000, failed to elicit natural competence (19), which may have been due too inappropriate expression levels of *comX*, or might have been caused by the fact that the endogenous *comX* gene of IL1403 was used which contains an alternative start codon and appears to be truncated.

Taken together, this study shows that in *L. lactis* strains that encode complete late-competence gene-sets, a state of competence can be induced by controlled expression of *comX*, in particular moderate expression of this regulator appears to be effective in activation of this phenotype. Naturally competent *L. lactis* strains could internalize plasmid and linear DNA from their environment with a similar efficiency. The conditions that naturally activate *comX* expression and contribute to the regulation of competence development in *L. lactis* remain to be established. Unraveling the *in situ* control mechanisms of natural competence in *L. lactis* would offer opportunities to exploit this phenotype for strain improvement purposes in this industrially important species.

## Materials & Methods

### Bacterial strains, plasmids, and media

The strains used in this study can be found in Table S1. The publicly available draft genome sequences of 43 *L. lactis* strains (20, 21, 25–34) were used for comparative genomics of late competence genes, by employing OrthoMCL to obtain orthologous group (OG) sequences in order to construct an orthologous gene matrix (61, Wels et al., manuscript in preparation). *L. lactis* strains were routinely cultivated in M17 (Tritium, Eindhoven, the Netherlands) supplemented with 1% (w/v) glucose (Tritium, Eindhoven, the Netherlands), at 30 °C without agitation. For competence experiments, *L. lactis* strains were cultivated in chemically defined medium (CDM) (62, 63) supplemented with 1% (w/v) glucose (Tritium, Eindhoven, the Netherlands). Upon recovery after electro or natural transformation, *L. lactis* cells were cultivated in recovery medium (M17, supplemented with 1% glucose, 200mM MgCl_2_ and 20mM CaCl_2_). *Escherichia coli* TOP10 (Invitrogen, Breda, The Netherlands) was routinely cultivated in TY (Tritium, Eindhoven, the Netherlands) at 30 °C with agitation. Antibiotics were added when appropriate: 5.0 μg/ml chloramphenicol, 10 μg/ml erythromycin, 12.5 μg/ml tetracyclin.

### DNA manipulations

Plasmid DNA from *E. coli* and *L. lactis* was isolated using a Jetstar 2.0 maxiprep kit (ITK Diagnostics bv, Uithoorn, The Netherlands). Notably, phenol chloroform extraction was performed prior to loading on the Jetstar column for plasmid isolation from *L. lactis* cultures (64). Primers were synthesized by Sigma-Aldrich (Zwijndrecht, The Netherlands). PCR was performed using KOD polymerase according to the manufacturer’s instructions (Merck Millipore, Amsterdam, The Netherlands). PCR products and DNA fragments in agarose gel were purified using the Wizard^R^ SV gel and PCR Clean-Up System (Promega, Leiden, The Netherlands). PCR-grade chromosomal DNA was isolated by using InstaGene™ Matrix (Bio-rad, Veenendaal, The Netherlands). Ligations were performed using T4 ligase and when applicable either transformed to electrocompetent *E. coli* TOP10 (Invitrogen, Breda, The Netherlands) or *L. lactis* NZ9000, IL9000, KW2 or KF147 (65).

### Plasmid and mutant construction

To enable controlled expression of *comX* in *L. lactis*, the *comX* gene was amplified by PCR using the primer pairs C1-C2 or C3-C4, and *L. lactis* KF147 or MG1363 chromosomal DNA as a template, respectively. The resulting 502 bp *comX* amplicons were digested with KpnI (introduced in primers C2 and C4) and ligated into KpnI-ScaI digested pNZ8150 (40), yielding pNZ6200 and pNZ6201, respectively. These *comX* overexpression vectors were transformed into electrocompetent *L. lactis* KF147, NZ9000, IL9000 and KW2 (65). The natural competence potential in *L. lactis* strains was evaluated using pIL253 when possible but because of incompatibility of antibiotic resistance markers in strain KW2 an alternative plasmid was constructed in which the erythromycin resistance (*eryR*) gene was replaced by tetracyclin resistance (*tetR*) gene. To this end, a 1644 bp *tetR* amplicon was generated using primers C15 and C16 with pGhost8 (66) as a template, and cloned as a PstI-SacI fragment into similarly digested pIL253, yielding pNZ6202.

A *comEA-EC* deletion derivative of *L. lactis* KF147 mutant was constructed using double cross over recombination. To construct the mutagenesis fragment, the 5’- and 3’- flanking regions of the *comEA-EC* operon were amplified using chromosomal DNA of strain KF147 as a template and primer pairs C7-C8, and C11-C12, respectively. The tetracylin resistance encoding gene *tetR* was amplified from pNZ7103 (67) using primers C9 and C10. SOEing PCR (68) was employed to join the three amplicons using the compatible sequence overhangs introduced by the primers in the individual PCR reactions (Table S1), and primers C7 and C12 for amplification in this PCR. The 6 kb SOEing amplicon was purified from 1% agarose gels and transformed to naturally competent *L. lactis* KF147 (see results section). The anticipated *comEA-EC* deletion in the resulting derivatives of *L. lactis* KF147 yielding *L. lactis* NZ6200 was confirmed by PCR using C13 and C14 primers.

### Induction of competence in *L. lactis*

Cells harboring pNZ6200, pNZ6201 or pNZ8150 were grown overnight in GCDM with appropriate antibiotics, followed by subculturing (1:65) in the same medium to an OD_600_ of 0.3 at which Ultrapure Nisin A (Handary, Brussels, Belgium) was added to the media at a final concentration of 0.005, 0.01, 0.03, 0.05, 0.07, 0.1, 0.5 or 2ng/ml. In parallel, 1 µg of plasmid DNA was added. Samples were incubated for 2h at 30°C, after which 5ml recovery medium was added and incubation was continued for another 2 h. Bacteria were pelleted by centrifugation at 4000xg for 8 minutes and transformants were enumerated by plating of serial dilutions on GM17 plates. KF147 Transformants were subjected to PCR analysis to assess presence of the transformed plasmid with primers PS1 and PS2, whereas the strain-specific primers SS1+2 were used to confirm strain identity.

### Analysis of competence gene expression

RNA was isolated from *L. lactis* cultures using the High Pure RNA isolation kit (Roche Diagnostics Nederland B.V., Almere, The Netherlands), including an on-column DNase treatment. Eluted RNA was again DNase (1 U; Thermo fisher scientific, Waltham, USA) treated for 45 min at room temperature to remove remaining DNA, followed by DNase inactivation by the addition of EDTA to a final concentration of 25mM, followed by heating at 75 °C for 15min. cDNA was prepared using 10 ng total RNA and random hexamer primers in the reverse transcriptase reaction (Applied Biosystems, Foster city, USA) according to the manufacturer’s protocol. Control RNA samples that were not reverse transcribed were included as negative control to ensure the absence of DNA contamination. Transcripts of competence genes were quantified using 2 μl cDNA and locus-specific primers for each competence associated target gene (Primers Q1-Q10, Table S1) in SYBR green-quantified PCR (Bio-rad, Veenendaal, The Netherlands). Transcript copy-numbers were calculated using a template standard curve and normalized to the housekeeping-control transcript of *rpoA*. These RT-qPCR analyses were performed in triplicate for each sample using the Freedom EVO 100 robot system (Tecan, Männedorf, Switzerland), and amplicon identities were verified using melting curve analysis. The non-parametric Mann-Whitney U-test (one-tailed) was used to determine whether gene expression levels were significantly different between uninduced and induced conditions (*P*-value<0.05).

## Acknowledgements

We gratefully acknowledge Prof. dr. Jan Kok for providing *L lactis* strain IL9000. We would like to thank Sabri Cebeci and Koen Giesbers of NIZO for technical assistance. This work was carried out within the BE-Basic R&D Program, which was granted a FES subsidy from the Dutch Ministry of Economic affairs.

